# Protein Compositional Ratio Representation (PCRR) Systematically Improves Human Disease Prediction

**DOI:** 10.1101/2025.11.29.691286

**Authors:** Adithya V. Madduri, Randall J. Ellis, Chirag J. Patel

## Abstract

Plasma proteomics captures a functional snapshot of human physiology; yet, most machine learning models treat protein abundances as independent variables, ignoring the fact that biological systems and proteomic measurements are inherently compositional. Many molecular processes depend not on absolute concentrations but on relative balances: receptor–ligand stoichiometry, enzyme–substrate ratios, and homeostatic feedbacks that govern signaling and metabolism. We propose that these relationships are best captured through pairwise protein ratios, which more faithfully reflect underlying biochemical constraints than raw expression values.

We evaluate a machine learning framework that models pairwise log-ratios of proteins (*log*(*A*) −*log*(*B*)) as features, thereby encoding compositional structure directly into the learning space. Applied to the ROSMAP plasma proteomics cohort (n = 871), this approach substantially improved the classification of Alzheimer’s subtypes (NCI, MCI, AD, AD+) with an average AUROC gain of +0.1274 over a strong baseline that incorporated raw proteomics and demographics. The top-ranked ratios (e.g., SEMA3C:TMEM70, IDUA:NPTXR) captured converging pathogenic pillars of Alzheimer’s disease, including microglial activation, proteostasis dysregulation, and lipid-clearance imbalance, highlighting that ratio-based features recover biologically coherent axes of disease.

To assess generality, we scaled the method to the UK Biobank proteomic dataset (n > 53,000; 587 phenotypes). The ratio-based model outperformed raw-level models in 95.1 % of diseases, with statistically significant (FDR < 0.05) gains in 56.7 %. Together, these results suggest that proteomic data should be viewed and modeled as compositional systems, where relative protein abundances carry the accurate functional signal. This insight supports the broader utility of ratio-based representations for disease prediction and biomarker discovery.

## 1 Introduction

High-dimensional proteomic data provide a powerful, dynamic window into human physiology and disease. As large-scale plasma profiling now enables the measurement of thousands of circulating proteins per individual, the challenge has shifted from data acquisition to representation. Models trained on absolute protein abundances often struggle with batch effects, normalization artifacts, and inter-individual variability, obscuring biological structure and limiting predictive power.

We hypothesize that the fundamental unit of proteomic variation is not the absolute level of a single protein, but the *relative balance* between proteins within shared pathways or regulatory systems. Biological processes such as receptor–ligand signaling, enzyme–substrate interactions, and feedback regulation are inherently compositional: it is the ratio between components that determines functional outcomes. This principle, long recognized in compositional data analysis (1), suggests that pairwise protein ratios may provide a more stable and biologically meaningful representation of the proteome. Mathematically, this can be captured by simple log-ratio features of the form log(*A*) − log(*B*), which are variance-stabilized, scale-invariant, and directly encode biological relationships that raw protein levels may obscure.

We developed a generalizable machine-learning framework to systematically test this ratio-based representation across diseases, beginning with Alzheimer’s disease (AD) as a case study in biological heterogeneity. AD affects over 30 million people worldwide (2) and remains one of the most molecularly and clinically heterogeneous neurodegenerative disorders known (3). Proteomic measurements are uniquely suited to probing this heterogeneity, as they capture dynamic molecular changes rather than static genetic risk.

Using plasma proteomic data from the Religious Orders Study and Memory and Aging Project (ROSMAP) cohort (4), we trained models to classify four AD subtypes (NCI, MCI, AD, and AD+). Our ratio-based framework was evaluated against a cascade of increasingly strong baselines: (1) a stratified random classifier, (2) a model using demographic covariates (age, sex, education, *APOE*), and (3) a model combining demographics with raw proteomic levels. Across all subtypes, models using only engineered log-ratios achieved an average AUROC improvement of +0.2525 over the random baseline, +0.1540 over the demographics-only model, and critically, +0.1274 even relative to the gold-standard baseline combining raw proteomics and demographics. Feature importance analysis of top-ranked ratios revealed processes central to AD pathology, highlighting that predictive ratios are also mechanistically interpretable.

To test whether this advantage reflects a broader principle of biological compositionality rather than an AD-specific effect, we extended our framework to the UK Biobank (UKB) (5). Analyzing proteomic data from over 53,000 individuals, we applied the same ratio-based feature engineering pipeline to predict 587 distinct disease outcomes spanning neurological, immune, metabolic, and other categories. The results were strongly concordant: ratio-based models out-performed models using raw protein levels in 95.1% of all outcomes, with statistically significant gains (FDR < 0.05) in 56.7%.

Together, these results reveal that plasma proteomic data are fundamentally *compositional* : the information that distinguishes disease states lies in the relative balance among proteins rather than their absolute abundances. This insight provides a unifying principle for proteomic data representation, offering a biologically grounded, interpretable, and general-purpose strategy for biomarker discovery and disease prediction.

## 2 Methods

Our methodological approach is divided into two parts. First, we develop and apply our ratio-based feature engineering pipeline to a deep, longitudinal cohort to classify heterogeneous Alzheimer’s disease subtypes. Second, to demonstrate the generalizability of this method, we apply the same pipeline at scale to an independent, cross-sectional cohort (the UK Biobank) across 587 distinct disease outcomes.

### 2.1 Compositionally-Aware Ratio Representation

Proteomic measurements are naturally viewed as compositional: only the *relative* abundances of proteins are biologically meaningful, while global scaling (for example due to technical factors or sample dilution) does not change the underlying state. Formally, let 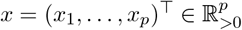 denote the strictly positive plasma abundances of *p* proteins. Two vectors *x* and *cx* for any scalar *c* > 0 represent the same composition, since they differ only by a global multiplicative factor. The set of equivalence classes 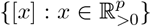 under this relation can be identified with the Aitchison simplex 𝒮^*p*−1^, the standard sample space for compositional data (1).

A classical approach in compositional data analysis is to map the simplex into a (*p* − 1)-dimensional real vector space using log-ratio transformations (1). Let *y*_*i*_ = log *x*_*i*_ for *i* = 1, …, *p* (any logarithm base is valid; in our experiments, we use base-10 and base-2 depending on the preprocessed data). The centered log-ratio (clr) transformation is defined as

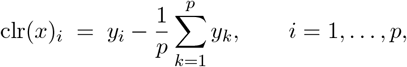

which embeds 𝒮^*p*−1^ into the (*p* − 1)-dimensional subspace {*z* ∈ ℝ^*p*^ : ∑_*i*_ *z*_*i*_ = 0}. Any classifier that depends only on clr coordinates is therefore invariant to global rescaling of the original abundances.

In this study, we do not use the clr transformation directly; instead, we construct pairwise log-ratios of the form:

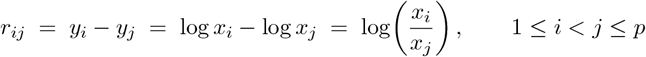

Collecting all such differences yields a linear transformation *r* = *Ay*, where *A* is a fixed matrix with entries in {−1, 0, 1}. Because adding a constant to all coordinates of *y* leaves every difference *y*_*i*_ − *y*_*j*_ unchanged, *r* depends only on the composition [*x*] ∈ 𝒮^*p*−1^ and not on its overall scale. Moreover, the space spanned by these pairwise log-ratios is the same clr subspace: each clr coordinate can be written as a linear combination of pairwise differences, and conversely each pairwise difference depends only on clr coordinates. Thus, the full set of pairwise log-ratios forms an overcomplete linear representation of the underlying composition, preserving all information up to a global multiplicative constant.

In practice, we do not enumerate all 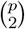 ratios. Instead, we first identify a data-driven short-list of consistently predictive proteins (Section 2.3.2), and then generate pairwise log-ratios only within this set. Conceptually, this corresponds to restricting the compositional representation to a lower-dimensional subset of proteins that carry most of the task-relevant signal, while still operating in a log-ratio space that respects the geometry of the simplex.

### 2.2 Protein Feature Representation

In this study, references to *raw, non-compositional*, or *non-ratio* protein features denote models trained on individual normalized protein abundance measurements, rather than on ratio-based representations. Importantly, these features are *not* unprocessed assay outputs. All protein measurements were subjected to standard dataset-specific normalization prior to modeling (e.g., log-transformation for ROSMAP and platform-specific normalization for UK Biobank). The distinction between models is therefore purely representational: non-ratio models treat normalized protein abundances as unconstrained coordinates in Euclidean space, whereas ratio-based models operate on log-ratios that respect the compositional geometry of the data. As a result, all comparisons isolate the effect of feature representation rather than differences in preprocessing or data quality.

#### 2.2.1 Scale invariance of the ratio representation

A direct consequence of the log-ratio construction is invariance to global multiplicative scaling: **Proposition 1**. Let 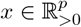 and *c* > 0. For any pair of indices *i, j* ∈ {1, …, *p*},

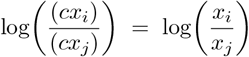

In particular, the pairwise log-ratio feature vector *r*(*x*) is identical for *x* and *cx*, and any classifier that operates solely on *r*(*x*) cannot be affected by global changes in assay scale or sample dilution.

This property formalizes the intuition that our Protein Compositional Ratio Representation (PCRR) removes irrelevant multiplicative noise and constrains learning to contrasts that are interpretable on the Aitchison simplex. The empirical results in Sections 3.1 and 3.2 show that, in both ROSMAP and UK Biobank, this compositional representation is substantially more predictive than treating raw protein levels as unconstrained coordinates in Euclidean space.

From an information-geometric standpoint, the Aitchison simplex is a nonlinear sample space whose intrinsic geometry reflects changes in relative proportions rather than absolute scale. By mapping compositions into log-ratio coordinates, the data are placed in a flat Euclidean subspace equipped with the Aitchison inner product, so that pairwise differences log(*x*_*i*_*/x*_*j*_) correspond to valid geodesic directions on this manifold. This provides a geometric interpretation for why ratio-based representations preserve the statistically meaningful structure of the composition while raw levels do not.

### 2.3 Alzheimer’s Disease Subtype Modeling (ROSMAP)

#### 2.3.1 ROSMAP Cohort and Phenotypes

We used the proteomic cohort from the Religious Orders Study and Memory and Aging Project (ROSMAP) (4), which consists of *n* = 871 individuals with *n*_*visits* = 953. The dataset comprises plasma proteomics data for 7,298 proteins, collected using the SomaScan assay. As previously described (6), the data was log-10 normalized due to its log-normal distribution, with each patient visit forming a feature vector in ℝ^7298^.Phenotypes were clinically determined at the time of each visit based on established clinical decision rules (7). We modeled four distinct classes: (1) No Cognitive Impairment (NCI); (2) Mild Cognitive Impairment (MCI); (3) Alzheimer’s Disease (AD); and (4) AD+ (indicating AD with a concurrent cause of cognitive decline).

#### 2.3.2 Ratio-based Feature Engineering for AD Subtypes

To model interactions and relative protein expression levels, we implemented a three-stage feature engineering pipeline to generate predictive pairwise ratios.

1. **Initial Feature Prioritization**: We first trained a multi-class LightGBM (LGBM) model on the full raw feature set (all 7,298 proteins plus demographics) across 5 random splits. A protein was considered “consistently predictive” if it exhibited non-zero feature importance in at least 2 of the 5 splits for any AD subtype class.
2. **Log-Ratio Generation:** From the resulting shortlist of *k* predictive proteins, we generated all unique, non-redundant pairwise log-ratios. For any two log-normalized proteins *P*_*a*_ and *P*_*b*_ from the shortlist (*a* < *b*), we computed their difference:

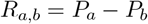

Since the input data was already log-10 normalized (e.g., *P*_*a*_ = log_10_(Protein A abundance)), this subtraction is mathematically equivalent to the log of the ratio of the original, non-normalized abundances: *R*_*a,b*_ = log_10_(*A*) − log_10_(*B*) = log_10_(*A/B*). This created a new feature matrix with 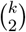 potential log-ratio features. Because both cohorts provide log-transformed protein measurements (ROSMAP: log_10_; UKB NPX: log_2_), ratio features were computed via subtraction in log space rather than explicit division, and we did not encounter numerical instability from small denominators.

The final “ratio models” for the ROSMAP cohort were then trained and evaluated using all engineered ratio features.

### 2.4 Generalizability Analysis (UK Biobank)

#### 2.4.1 UK Biobank Cohort and Disease Outcomes

To test the broader utility of our method, we applied it to the UK Biobank (UKB) cohort. This independent dataset consists of *n* > 53, 000 individuals with plasma proteomics data (Olink platform, 3,000 proteins). We analyzed 587 distinct disease outcomes, defined by ICD-10 codes, spanning a wide range of neurological, metabolic, immune, and other disease categories.

#### 2.4.2 UKB Feature Engineering Pipeline

We applied a modified version of our feature engineering pipeline to accommodate the scale of the UKB analysis. For each of the 587 disease outcomes, we performed the first two steps of our pipeline: (1) Initial Feature Prioritization (training on raw proteins to find “consistently predictive” proteins) and (2) Ratio Generation (creating all pairwise ratios from the predictive shortlist). Importantly, the shortlist of proteins was derived exclusively from training folds within the cross-validation of the initial protein-level models; no test labels or outcome information were used in feature selection. As a result, the subsequent ratio models, trained on the same fold structure, used only training-derived features, ensuring that no information leakage occurred. Models were trained on the full set of 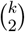 generated ratios to assess the raw predictive power of the ratio-based feature space

### 2.5 Model Training, Validation, and Evaluation

#### 2.5.1 Experimental Design and Validation

All classifiers were trained in a one-vs-rest (OVR) setting for multi-class problems.

1. **For ROSMAP**, we used a repeated hold-out validation strategy across five random 70/30 (train/test) splits. To prevent data leakage from the longitudinal data, these splits were group-aware, ensuring all visits from the same patient were assigned exclusively to either the training or test set.
2. **For UK Biobank**, we used a standard 5-fold cross-validation for each of the 587 disease endpoints.

In both cohorts, splits were stratified by the target class label and sex to maintain consistent class distributions.

#### 2.5.2 Model Architectures and Hyperparameter Optimization

For the initial ROSMAP cohort analysis, we evaluated two interpretable, tree-based ensemble models: **Random Forest (RF)** (8) and **LightGBM (LGBM)** (9). We used A Fast Library for Automated Machine Learning & Tuning (FLAML) (10) to optimize hyperparameters for both architectures. In these head-to-head comparisons for AD subtype classification, LGBM models consistently demonstrated superior predictive performance over RF. Based on this finding, we selected **LGBM as the sole architecture** for the large-scale UKB generalizability analysis to ensure computational efficiency while using the best-performing model. For all experiments (both RF and LGBM on ROSMAP, and LGBM on UKB), hyperparameter optimization was conducted via FLAML. For each of the five evaluation splits, we allocated FLAML a time budget of 100 seconds per class (a fixed, extended budget of 1200 seconds was used for ratio models to account for increased feature space size), which used 5-fold internal cross-validation on the training set to find the optimal hyperparameters. RF models were implemented via scikit-learn (11) and LGBM via the official LightGBM package.

#### 2.5.3 Evaluation and Baselines

Model performance was primarily assessed by the Area Under the Receiver Operating Characteristic Curve (AUROC) and Average Precision (AP), with results averaged across the evaluation splits.To establish a performance floor and contextualize our results, we compared our proteomic ratio models against several baselines:

1. **Stratified Random:** A chance-level classifier that predicts by sampling from the training set’s class distribution.
2. **Demographics-Only:** A model trained on standard clinical covariates: age, sex, years of education, and *APOE* genotype (or equivalent available covariates for UKB).
3. **Raw Proteomics + Demos:** The strongest baseline, trained on the full set of 7,298 raw (log-10 normalized) protein levels in addition to all demographic covariates.

## 3 Results

### 3.1 Proteomic Ratio Model Improves AD Subtype Classification

To determine whether proteomic ratios could improve prediction of disease outcomes, we began by evaluating our log-ratio feature engineering pipeline on the ROSMAP cohort to classify four challenging, clinically-defined AD subtypes: NCI, MCI, AD, and AD+. To rigorously quantify the benefit of our method, we compared the performance of our **Log-Ratio Model** against a cascade of three increasingly stringent baselines.

#### 3.1.1 Ratios Outperform Raw Data Plus Demographics

Our primary analysis used the Area Under the Receiver Operating Characteristic Curve (AU-ROC) to evaluate the predictive accuracy of model’s incorporating different data modalities. The baseline models revealed the difficulty of the classifying AD subtypes. The **Stratified Random** classifier performed at chance, with mean AUROCs of 0.5195 for MCI, 0.4890 for NCI, 0.4896 for AD, and 0.5387 for AD+ (Table 1). The **Demographics-Only** model showed modest predictive power, achieving AUROCs of 0.5508 for MCI, 0.6447 for NCI, 0.6137 for AD, and 0.6218 for AD+. Our strongest baseline, the **Raw Proteomics + Demos** model, improved on this, setting a robust performance benchmark with AUROCs of 0.5176 for MCI, 0.6749 for NCI, 0.7161 for AD, and 0.6288 for AD+ (Fig. 1a, Table 1).

**Figure 1.**
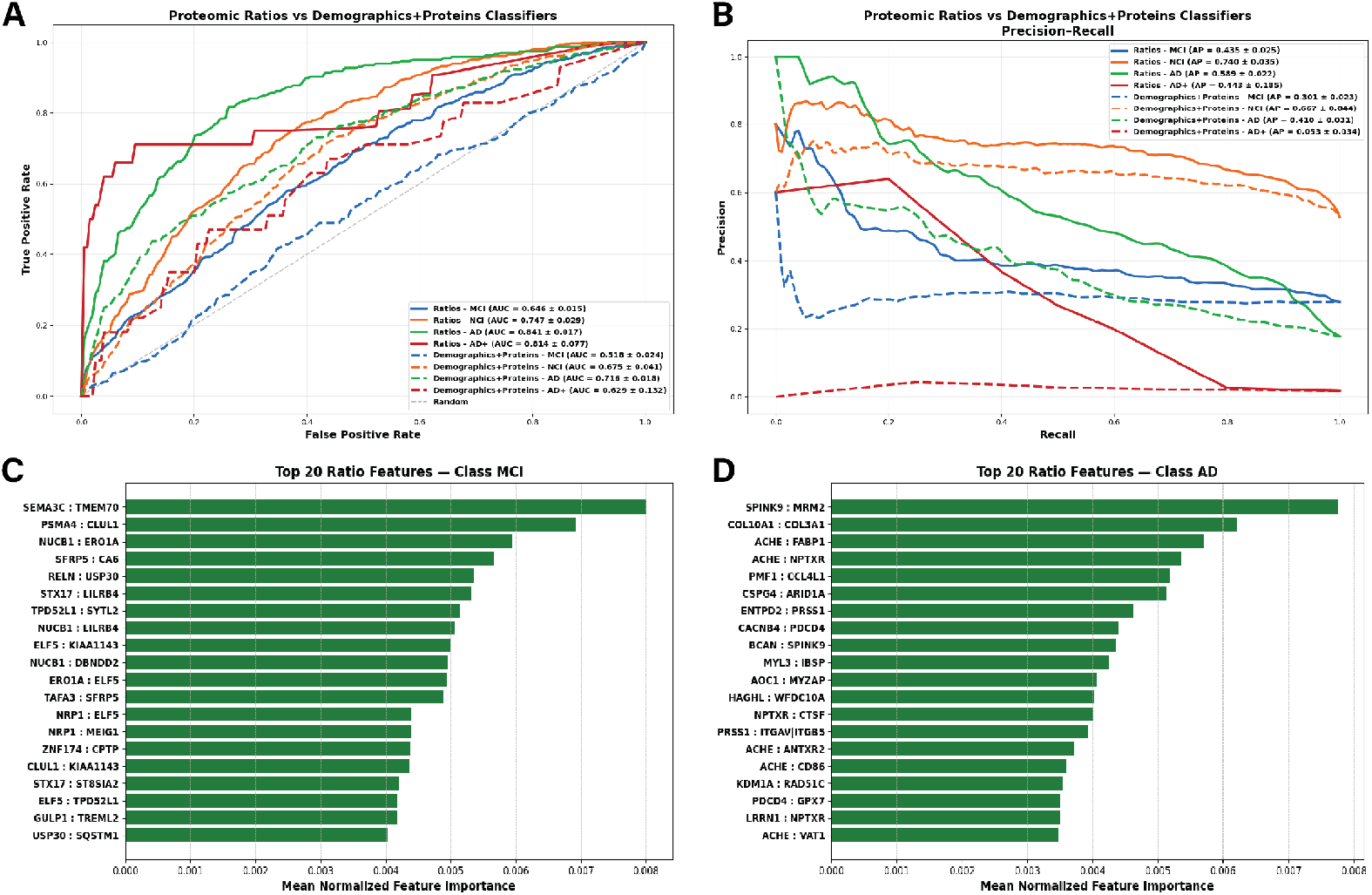
Overview of ROSMAP Cohort Results. a) ROC curve performance for the ratio based model (solid line) and demographics plus proteins model (dashedline), b) Precision-Recall curves for the ratio based model (solid line) and demographics plus proteins model (dashedline), Feature importance analysis for the Mild Cognitive Impairment subset of ROSMAP cohort, Feature importance analysis for the Alzheimer’s Disease subset of ROSMAP cohort.

Remarkably, our ratio-based Model, using just pairwise log ratios and no demographic data, consistently and substantially outperformed all baselines across every class (Fig. 1a). Our model achieved an AUROC of 0.6457 for MCI, 0.7469 for NCI, 0.8407 for AD, and 0.8136 for AD+. Most critically, even when compared to the strongest **Raw Proteomics + Demos** baseline, our log-ratio model achieved an average AUROC improvement of +0.1274 across all classes. This demonstrates that our feature engineering method isolates a more predictive signal than the raw data alone. The gains were largest for the most difficult-to-classify subtypes, with a +0.1281 AUROC improvement for MCI and +0.1848 for AD+, along with gains of +0.0721 for NCI and +0.1246 for AD (Table 1). When compared to the **Demographics-Only** model, the log-ratio features yielded an even greater average AUROC gain of +0.1540, with class-specific improvements of +0.0949 (MCI), +0.1022 (NCI), +0.2270 (AD), and +0.1918 (AD+). Finally, as expected, the model showed massive gains over the chance-level **Stratified Random** baseline (average improvement of +0.2525), most notably with a +0.3511 AUROC improvement in AD classification (Fig. 1a). On average, across all three baselines our log-ratio model showed an improvement of +0.1780 (Table 1).

#### 3.1.2 Average Precision Analysis: Gains are Magnified in Imbalanced Classes

We next evaluated performance using the Average Precision (AP) score, a more informative metric for imbalanced clinical data where the superiority of the log-ratio method was even more dramatic. The **Stratified Random** model APs were 0.3067 (MCI), 0.5310 (NCI), 0.1856 (AD), and 0.0652 (AD+). The **Demographics-Only** model offered little improvement for minority classes, with APs of 0.3349 (MCI), 0.6555 (NCI), 0.2592 (AD), and a near-zero 0.0332 (AD+) (Table 1). The **Raw Proteomics + Demos** model, our strongest baseline, struggled significantly with the minority AD+ class, achieving an AP of just 0.0531, along with 0.3011 (MCI), 0.6670 (NCI), and 0.4102 (AD) (Table 1). This indicates the baseline models almost completely failed to reliably identify the AD+ class.

In stark contrast, our **Ratio-based model** achieved a robust AP of 0.4348 for MCI, 0.7398 for NCI, 0.5887 for AD, and 0.4429 for AD+, demonstrating a strong ability to identify all classes (Fig. 1b, Table 1). These improvements were substantial across all comparisons. Our method yielded an average AP improvement of +0.1937 over the **Raw Proteomics + Demos** baseline (Fig. 1b). The most staggering gain was seen in the AD+ class, which improved by +0.3898 (a more than 8-fold increase in AP), effectively turning an unusable classifier into a viable one. We also saw major gains for MCI (+0.1337), AD (+0.1785), and NCI (+0.0727). Against the **Demographics-Only** model, the average AP improvement was +0.2308, with class-specific gains as high as +0.4097 for AD+ and +0.3295 for AD. Finally, relative to the **Stratified Random** baseline, our model achieved an average AP improvement of +0.2794, including a +0.4031 improvement for the AD class and +0.3777 for the AD+ class (Table 1).

Taken together, these results across both AUROC and AP metrics show that our ratio feature engineering pipeline is not just additive, but transformative. It isolates highly predictive, biologically-relevant interactions that are otherwise obscured in raw proteomic data, enabling superior and more robust classification of heterogeneous AD subtypes, especially for difficult minority classes.

#### 3.1.3 Feature Importance Profiles Across AD Subtypes

We next examined the most influential ratio features contributing to classification within each diagnostic category. Given the improvement in predictive accuracy achieved by PCRR, these feature importance profiles provide a principled way to identify disease-relevant protein contrasts.

In the MCI model, ratios involving SFRP5 (an anti-inflammatory protein) (12), TMEM70 (reported to be downregulated in AD), and NUCB1 emerged as the most recurrent and stable predictive features, with the SEMA3C:TMEM70 ratio ranking as the single most informative contrast (Fig. 1c). TREML2 also appeared prominently among the top predictors, a notable finding given the extensive literature linking TREM2 receptor expression levels to microglial detection of Amyloid and subsequent adoption of the Disease Associated Microglia (DAM) state in the presence of Alzheimer’s pathology (13). Additional consistently selected predictors included TPD52L1 and TMEM70, further highlighting the convergence of ratio-based signals on proteins implicated in cellular stress, immune activation, and early neurodegenerative processes. CLUL1 was another notable predictor, consistent with CLU as a repeated GWAS hit in AD and its broader roles in immune regulation and amyloid-*β* aggregation (14).

In the AD model, the top ratios prominently involved SPINK9, MRM2, COL10A1, COL3A1 and multiple ACHE-based contrasts (Fig. 1d). Notably, acetylcholinesterase (ACHE) has been implicated in AD pathology, where it not only modulates cholinergic signaling but may also interact with amyloid-*β* and influence plaque formation, consistent with its prominence in multiple top ratio features (15). ARID1A was also repeatedly implicated, in alignment with prior literature linking ARID1A to AD (16).

Across both NCI and AD+, we identified ratios involving proteins previously associated with AD as well as potentially novel ratio-based biomarkers. Notably, many of these proteins have been studied largely in isolation, whereas our framework nominates informative ratios that combine both well-established AD drivers and less-characterized proteins, highlighting interactions that may be overlooked by single-protein analyses.

### 3.2 Generalizability Analysis

To assess the generalizability of our framework, we evaluated the ratio-based approach across 587 disease outcomes in the UK Biobank. The ratio-based model outperformed raw proteomics in 558 outcomes (95.1%), with significant improvements observed in 333 outcomes (56.7%) (Fig. 2c). Only 4 outcomes (0.7%) showed significantly worse performance. Overall, the ratio-based representation increased AUC by an average of 7.93%, with a maximum observed improvement of 46.6% (Fig. 2b).

**Figure 2.**
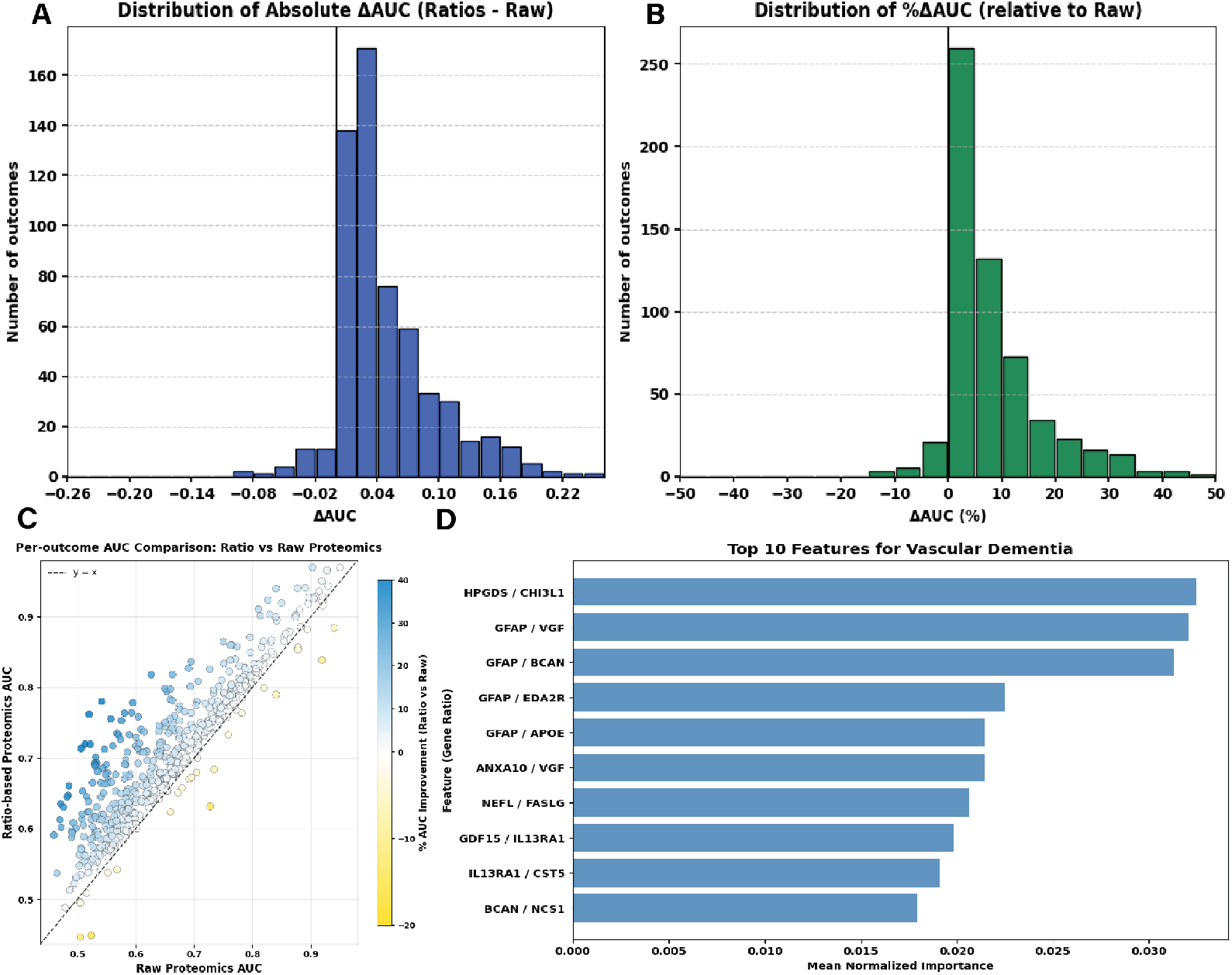
Overview of UKB Generalizability Analysis. a) Distribution of absolute AUC change between raw and ratio model, b) Percentage AUC change, c) Raw versus Ratio AUC per outcome, d) Top 10 features for Vascular Dementia.

#### 3.2.1 Discriminative Performance Across UK Biobank

Across these UK Biobank disease outcomes, the ratio-based representation yielded a mean AU-ROC improvement of 7.93% (median 5.05%) relative to raw protein models, with absolute changes ranging from a small minority of modest decreases (minimum Δ AUROC −0.096) to substantial gains (maximum Δ AUROC +0.242), indicating that while losses are limited, the potential upside is large. Consistent with this shift, the number of outcomes exceeding stringent absolute-performance thresholds increased markedly under PCRR, rising from 30 (5.11%) to 47 (8.01%) for AUROC ≥ 0.85 and from 288 (49.06%) to 397 (67.63%) for AUROC ≥ 0.65 when comparing raw protein models to ratio-based models.

#### 3.2.2 Outcome Interpretation

Several clinically important and biologically complex conditions showed clear improvements when using proteomic ratios, including Parkinson’s disease, vascular dementia, schizophrenia, hereditary ataxia, dystonia, and other degenerative and demyelinating diseases of the central nervous system. Beyond neurological conditions, strong gains also appeared across major cardiometabolic diseases such as acute myocardial infarction, chronic ischaemic heart disease, heart failure, atrial fibrillation and flutter, cerebral infarction, intracerebral haemorrhage, stroke, chronic obstructive pulmonary disease, asthma, chronic renal failure, obesity, iron-deficiency anaemia, other anaemias, and unspecified diabetes mellitus. These phenotypes represent large disease burdens with long preclinical windows and systemic metabolic, immune, or endocrine components, making them especially meaningful contexts in which proteomic ratios can plausibly capture early physiological changes that raw protein levels do not.

A large number of acute infectious diseases also showed significant improvements, including measles, varicella (chickenpox), rubella, viral meningitis, acute poliomyelitis, acute hepatitis A, acute hepatitis B, and a wide range of acute upper-respiratory, gastrointestinal, skin, and ear infections. Since the UKB proteomics measurements are prospective for most participants, these signals are unlikely to reflect pathogen-specific prediction years before the event. Instead, they most plausibly index baseline host vulnerability, including chronic inflammatory tone, immune competence, metabolic status, or other factors shaping susceptibility to future acute infections. This pattern is consistent with the observation that proteomic ratios enhance the detection of broad systemic states, rather than predicting the timing of specific infectious episodes, and suggests that ratio-based representations may be particularly sensitive to underlying biological frailty and immune predisposition.

#### 3.2.3 Feature Importance Analysis

As initial case studies, we first examined vascular dementia and Parkinson’s disease (PD) because their molecular signatures are well characterized, making it a highly interpretable outcome for evaluating feature coherence. Several of the highest-ranking ratios prominently featured GFAP, including GFAP:VGF, GFAP:BCAN, GFAP:EDA2R, and GFAP:APOE, reflecting a consistent astrocytic-injury signal across folds. Additional top ratios such as NEFL:FASLG and GDF15:IL13RA1 included proteins with well-established connections to neuronal injury or inflammation (Fig. 2d). Importantly, the presence of GFAP, APOE, and NEFL within the leading ratios aligns well with existing dementia and neurodegeneration literature, suggesting that the ratio-based model is capable of detecting real biological signal (17; 18). PD ratios included combinations such as ELN:PRL, ITGAV:PRL, SCG2:NEFL, EDA2R:ITGAV, and PRL:SCG2, which highlight reproducible model preferences across folds but also map onto established and potentially novel PD biomarkers (19).

## 4 Discussion

In this study, we introduce a generalizable framework for modeling plasma proteomics through pairwise log-ratios, motivated by the hypothesis that proteomic information is fundamentally encoded in relative rather than absolute abundances. Across two independent cohorts—ROSMAP and the UK Biobank—we demonstrate that these compositional representations consistently and substantially outperform models trained on raw protein levels augmented with demographic covariates. In Alzheimer’s disease, ratio-based models improved subtype prediction by an average AUROC of +0.1274 over the strongest baseline while sharply increasing average precision in minority classes such as AD+. At population scale, the same approach yielded significant gains in 56.7% of 587 UKB disease outcomes, with improvements spanning neurological, cardiometabolic, immunologic, and infectious phenotypes.

The mathematical motivation for our approach follows directly from the fact that proteomic measurements are inherently compositional: relative differences between proteins are meaningful, while absolute scales fluctuate due to assay effects and multiplicative noise. When models treat raw protein levels as independent coordinates in Euclidean space, they implicitly assume unconstrained variation that proteomic data do not possess. In contrast, log-ratio features, implemented here as simple differences log(A)–log(B), map compositional data into a space where scale invariance and proportional relationships are correctly preserved. This transformation removes irrelevant multiplicative biases, stabilizes variance, and ensures that proteins are compared in a way that reflects how they actually co-vary biologically. In effect, pairwise logratios provide a representation that respects the true constraints of proteomic data and allows machine-learning models to learn from meaningful contrasts rather than from absolute levels distorted by noise or scaling.

Biologically, many cellular processes depend on the balance between proteins rather than their individual concentrations. Receptor–ligand signaling, enzyme–substrate dynamics, lipid trafficking, immune activation, and proteostasis all operate according to relative availability within shared pathways. When these systems are perturbed, it is often the shift in balance, not the absolute abundance, that carries functional significance. Across phenotypes, these ratios can reveal coordinated molecular shifts that raw levels fail to isolate, providing biologically interpretable axes of dysfunction that align with known disease biology. Unifying these mathematical and biological interpretations, here we offer a rigorous proof of principle that proteomic data can and should be modeled as compositional.

### 4.1 Limitations

Our study has several limitations. While PCRR consistently improves predictive performance across cohorts and phenotypes, it is not a universal replacement for absolute protein quantification, and its utility will depend on biological context, measurement modality, and downstream task. Furthermore, our models rely on pairwise ratios generated after initial feature prioritization, and future efforts could explore more systematic or theoretically grounded selection strategies, including sparse log-contrast formulations or graph-aware ratio construction. Additionally, while our approach improves predictive performance and interpretability, it does not by itself establish causal mechanisms; follow-up experimental or perturbational studies will be needed to validate the biological pathways suggested by top-ranked ratios. Our work focuses on tree-based classifiers commonly used in biomedical machine learning to isolate the effect of representation choice; certainly, many alternative feature constructions are possible (e.g., additive, multiplicative, or higher-order interaction features), but our goal here was not to exhaustively benchmark every transformation, and pairwise log-ratios provide a principled, compositional baseline. Additionally, because log-ratios are transitive, the resulting feature space may contain multicollinearity, so individual ratio importances should be interpreted as reflecting broader compositional axes rather than unique causal edges. The pairwise log-ratio construction represents a deliberate trade-off between expressivity and scalability, providing a practical and interpretable alternative to higher-order compositional balances. We focus on pairwise log-ratios as a practical sweet spot: they encode biologically interpretable balances while remaining scalable and model-agnostic.

## 5 Future work

Looking ahead, several directions naturally extend from this work. As additional large-scale proteomic datasets become available, particularly those spanning diverse ancestries, platforms, and clinical contexts, evaluating the stability and portability of ratio-based representations will be essential for understanding their generalizability. Another opportunity lies in testing whether the same compositional principles translate to other high-dimensional molecular layers such as RNA-seq, metabolomics, or lipidomics, where relative shifts rather than absolute levels may likewise encode the most meaningful biological variation. More broadly, integrating ratio-based features across multi-omic modalities may reveal higher-order compositional structure that single-layer analyses cannot capture, highlighting clear next steps for both methodological refinement and biological investigation of compositional modeling.

### 6 Appendix

#### 6.1 Data and Code Availability

The plasma proteomics data used in this study was collected as part of the ROSMAP study and is available at: https://www.synapse.org/Synapse:syn64957327. The data are available under controlled use conditions set by human privacy regulations. To access the data, a data use agreement is needed. This registration is in place solely to ensure anonymity of the ROSMAP study participants. A data use agreement can be agreed with either Rush University Medical Center (RUMC) or with SAGE, who maintains Synapse, and can be downloaded from their websites. The UK Biobank proteomic data were accessed under application number 52887. These data are available to researchers through the UK Biobank Access Management System (https://www.ukbiobank.ac.uk/register-apply/), subject to approval and a data use agreement in accordance with UK Biobank’s access policies. All code used in this study is available at: https://github.com/AdiVM/Protein_Ratio_Representation.

#### 6.2 Cohort-level Normalization

As PCRR is defined on log-scaled protein measurements, the normalization and preprocessing procedures applied by each cohort are essential context for interpreting results. Here we summarize the assay modality and normalization pipeline used in each data source, and clarify how these values enter our feature construction; further details can be found at the dataset resources provided above.

##### 6.2.1 UK Biobank (UKB) proteomics

UK Biobank plasma proteomics were generated using the Olink Explore 1536 platform. The released protein measurements are provided as NPX (Normalized Protein eXpression) values, which are on a log_2_ scale. NPX values are produced via Olink’s standard processing pipeline, which includes internal controls and dataset-level normalization procedures to mitigate systematic technical variation. In particular, the released NPX values incorporate plate- and batch-level normalization, as well as bridge-sample normalization to improve comparability across plates and batches. We use these dataset-provided NPX values directly as the input to all UKB models and do not apply additional normalization beyond standard missingness filtering and covariate preprocessing described elsewhere.

##### 6.2.2 ROSMAP proteomics

ROSMAP plasma proteomics were generated using the SomaLogic SomaScan 7k aptamer-based assay. The released ROSMAP protein measurements are provided as log-transformed values (log_10_ scale) following dataset-provided normalization and quality control. These steps include standard SomaScan processing procedures to reduce assay-specific artifacts and to harmonize measurements across samples within the cohort. We use the normalized and QC-filtered values distributed with the dataset and do not perform additional cohort-level normalization beyond the preprocessing steps described in the main Methods.

##### 6.2.3 Scope and Limitations

Both cohorts used in this study rely on affinity-based proteomic platforms (Olink Explore and SomaScan) rather than mass-spectrometry-based quantification. We therefore do not claim that PCRR’s performance improvements generalize to all proteomic modalities, including LC-MS/MS acquisition strategies such as DDA, DIA, or targeted MS. Evaluating PCRR under additional proteomic platforms and quantification regimes remains an important direction for future work. Note that because UKB NPX values are on a log_2_ scale and ROSMAP values are on a log_10_ scale, ratio features in each cohort are computed within the cohort’s native log base.

#### 6.3 UKB AUC Analysis

For each of 587 disease outcomes in UK Biobank, we computed the mean test AUROC across 5 cross-validation folds for a LightGBM model trained on raw protein NPX values (Fig 3a) and the same model trained on PCRR pairwise log-ratios (Fig 3b) constructed from the training-derived predictive protein shortlist. Histograms use identical binning and x-axis limits to enable direct visual comparison of absolute discriminative performance.

**Figure 3.**
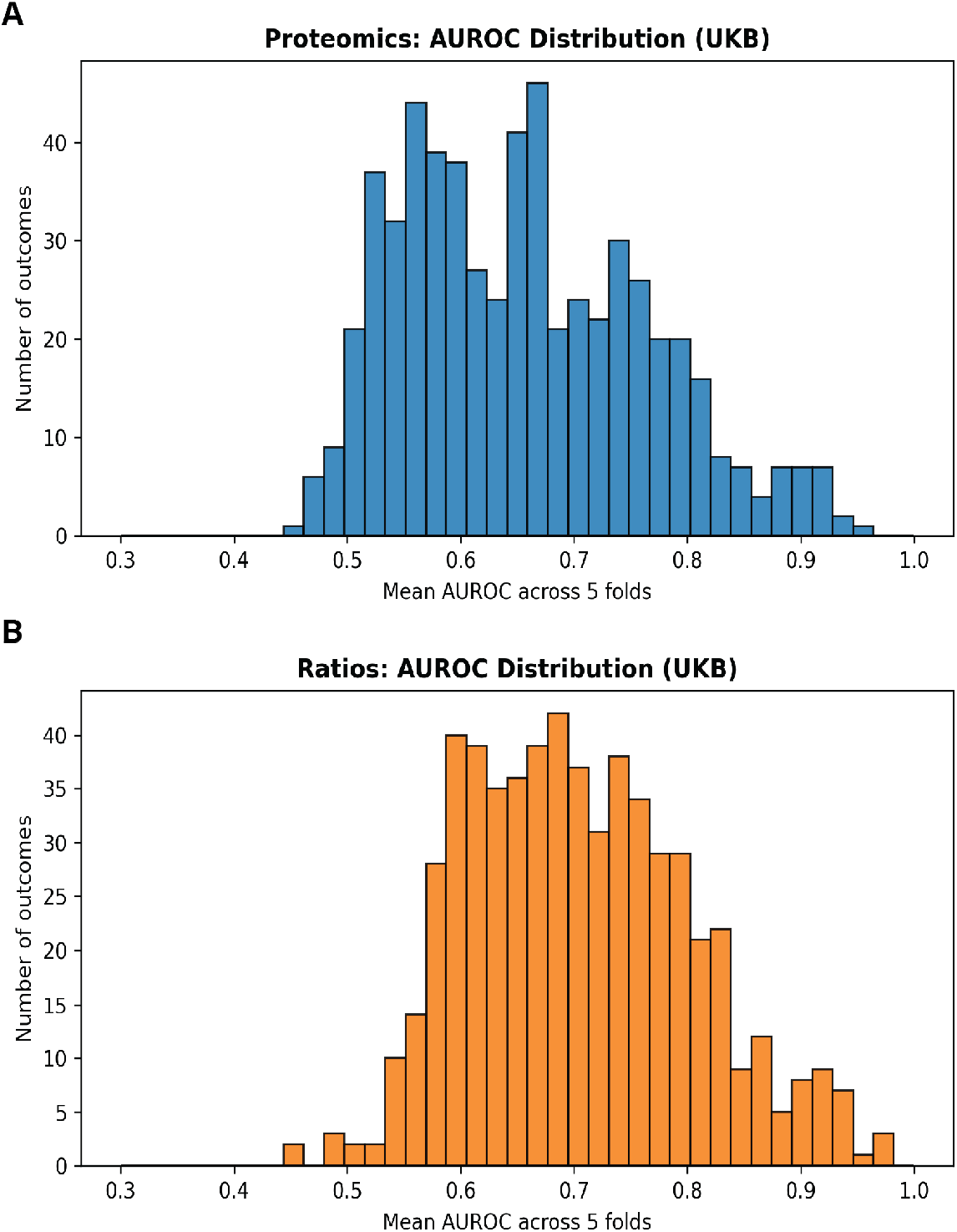
UK Biobank absolute AUROC distributions for raw proteins vs. PCRR.

#### 6.4 LASSO Generalization Analysis

To assess model independence, we tested whether PCRR’s benefit persists under an identical end-to-end pipeline by swapping only the base estimator from LightGBM to L1-regularized logistic regression. All preprocessing, data splits, and evaluation procedures, such as time budgets, were held fixed. Using the same preprocessing and fixed data splits, we trained an lrl1 baseline on the exact raw proteomics+demographics feature matrix used for the LGBM baseline, then generated log-ratio features from the same set of proteins identified by the LGBM feature-importance screen and trained an lrl1 classifier on these ratios. Across all four AD subtypes, the ratio representation consistently improved mean AUROC: 0.5679 to 0.6300(Δ = +0.0621) for MCI, 0.6288 to 0.7451(Δ = +0.1164) for NCI, 0.6368 to 0.8405(Δ = +0.2037) for AD, and 0.6872 to 0.8810(Δ = +0.1938) for AD+, indicating that the gains are not specific to tree-based models and persist under a linear classifier.

## Notes

### Competing Interest Statement

The authors have declared no competing interest.

### Summary of Updates

We have updated the abstract and title page.

